# The body size and fitness match and its variability in plastic response to temperature

**DOI:** 10.1101/2024.10.16.618639

**Authors:** Antoni Żygadło, Agata Burzawa, Katarzyna Potera, Franek Sierpowski, Aleksandra Walczyńska

## Abstract

The ability of organisms to respond plastically to environmental change remains one of the most challenging areas of ecological studies. Reasons for this include the complex nature of environmental cues and responses at the organismal level, the costs that organisms face in performing phenotypic plasticity under different conditions, and the individual capacity to respond, which depends on many internal and external factors. The special case in this regard is an example of the plastic body size response to temperature, known as the Temperature-Size Rule (TSR). We used five clonal and three multiclonal experimental populations of the *Lecane inermis* rotifer. We measured two traits over a wide thermal range: body size and population growth rate *r*. We investigated (i) the thermal conditions under which different experimental populations perform the TSR or canalize (= no plasticity) their body size, (ii) how this response relates to a direct measure of fitness, and (iii) whether this response varies with the thermal preferences of the studied organisms. We found a relationship between body size and fitness, confirming that the TSR is only performed within a certain thermal range, beyond which body size is canalized. We did not find the expected relationship between the strength of the TSR and the range of thermal tolerance, but our results do not allow us to reject the existence of such a relationship. Additionally, we found a relatively high repeatability of thermal tolerance in experimental populations in comparison with the previous studies. Finally, we suggest the role of the level of genetic composition within a population in its ability to respond plastically to temperature. This is the first study to describe in detail the special case of plasticity *versus* canalization, for body size response to optimal and suboptimal temperatures, in organisms that differ in their tolerance to temperature.

## 1. Introduction

Most organisms face varying degrees of heterogeneity in their native habitats. One possible strategy to overcome this challenge is phenotypic plasticity (DeWitt and Langerhans 2004, Schulte et al. 2011, Burraco et al. 2022), the ability of one genotype to express different phenotypes in different environments (West-Eberhard 1989). The adaptation of the phenotype to given environmental conditions through phenotypic plasticity represents the potential to evolve the trait in a longer time scale, as suggested by the ‘plasticity-first’ hypothesis (reviewed by Levis and Pfennig 2016). On a shorter time scale, adaptive phenotypic plasticity is the first-line organismal response to any environmental change. From this perspective, phenotypic plasticity plays a crucial role in organismal adjustment to environmental changes, including global warming, and understanding its mechanisms and limits is urgently needed (Fox et al. 2019).

The main benefit of phenotypic plasticity for a given genotype is that it increases the geometric mean fitness by reducing fitness variance between generations (Moran 1992). However, like any type of organismal response, phenotypic plasticity entails energetic costs. The general costs are associated with maintaining alternative phenotypes in different environments, while the direct metabolic costs are related to maintaining the organismal apparatus to perceive the environmental changes and adjust physiology accordingly (DeWitt et al. 1998, Relyea 2002). As life history strategies differ between organisms (Roff 1992, Stearns 1992, Kozłowski 2006), so do the costs associated with certain mechanisms, including those related to phenotypic plasticity. The evolution of phenotypic plasticity is an issue that we need to become familiar with in order to understand how environmental changes and disturbances affect communities (Auld et al. 2015, Gibert et al. 2019).

One of the tools that can be used to compare patterns of phenotypic plasticity between different organisms is the use of thermal performance curves (TPCs). These curves provide information on the general performance of the ectotherm in response to temperature changes, in particular the thermal tolerance range for a particular trait under investigation and the thermal optimum at which that trait reaches its maximum value (Huey and Kingsolver 1989). In its standard form, this is an asymmetric curve with a single maximum, skewed towards low temperatures (Huey and Kingsolver 1989). The asymmetric shape of the TPCs is mainly due to the fact that the measured traits describing performance and/or fitness are quantitative, with a complex dependence on temperature (Kingsolver 2001). The simplest patterns of TPCs distinguish organisms that prefer colder *vs*. hotter environments, have ’faster’ *vs*. ’slower’ lifestyles, or are generalists *vs*. specialists, *sensu* Levins (1968) (Kingsolver 2001). Other reasons for the shape of TPCs to vary include the time scale of a study, the thermal variability being studied, or the underlying trade-offs between different biochemical pathways (Schulte et al. 2011). Fitting such complex curves is not a trivial task. Recently, Padfield et al. (2021) provided a method for fitting 24 different TPC models, implemented in R language. Studies on the variability of TPCs between closely related organisms are relatively rare (Montagnes et al. 2021). For example, experimental studies on sepsid flies show that TPCs differ between closely related species, and between populations of the same species originating from different geographical regions (Khelifa et al. 2019). Further studies of this type are needed to determine how evolutionarily conserved TPCs are (Montagnes et al. 2021) and how much they may be influenced by the thermal history of the studied organisms (Pallares et al. 2021).

One of the most important responses to warming observed in communities on a global scale is reduced body size (Daufresne et al. 2009, Martins et al. 2023). Such a pattern could be caused by several factors, but one may be the plastic response of smaller body size at maturity with higher temperature, observed in over 80% of ectotherms, including unicellulars, plants and animals, known as the temperature-size rule (TSR; Atkinson 1994). There are many uncertainties about this rule, one of which is how and why the strength of the TSR response varies between groups of organisms. There are many debated ideas about the causes of variability in TSR strength, as reviewed by Verberk et al. (2021), but definitive conclusions remain elusive. In his classic work on the performance of organisms exposed to heterogeneous environments, Levins (1968) distinguished the two strategies, generalists, those that perform relatively well in all types of environments, and specialists, those that perform better around their environmental optimum. A link between the strength of the TSR and the organismal strategy of specialist or generalist was first suggested for the *Brachionus plicatilis* (Rotifera) cryptic species (Walczyńska and Serra 2014a) and confirmed at the interclonal level for another rotifer species, *Lecane inermis* (Stuczyńska et al. 2021). This latter study found an interrelated relationship between thermal preference, body size and the strength of the TSR. Specifically, generalists showed a stronger body size response to changing temperature (= larger absolute slope of body size change with temperature) than specialists. This topic is important in the light of a substantial debate about the differences in vulnerability to climate change between generalists and specialists (e.g., Huey et al. 2012), and their implications at the community-level response (e.g., Ohlberger 2013).

Instead of showing the TPC shape, body size decreases with increasing temperature. Nevertheless, this pattern does not refer to the whole thermal range. The existence of a specific thermal zone within which the TSR is performed was first conceptualised by Atkinson et al. (2003). Later, Walczyńska et al. (2016) conducted tests for the boundaries for the TSR performance in three species representing a unicellular (protist), eutelic multicellular (rotifer) and non-eutelic multicellular (annelid). The results revealed for all investigated organisms a body size decrease with increasing temperature only within a certain thermal range, restricted between temperature at which population growth rate approached zero (thermal minimum; T_min_), and temperature at which it reached the maximum value (thermal optimum temperature; T_opt_). Interestingly, beyond this range, body size was canalized (no plasticity; sensu Waddington 1942) and this canalization caused an intriguing reverse pattern for plastic body size changes around the T_min_ and T_opt_ (Fig. 4 in Walczyńska et al. 2016). The idea of canalization of organismal traits against genetic and environmental perturbations is not new, but it is still intriguing and not fully resolved (Schneider 2022). Stearns and Kawecki (1994) stated that any mechanisms that canalize a trait to make it closer to optimum should be favoured by natural selection. Both, genetic and environmental canalization evolved to maintain the stable reaction norm across environments (Stearns and Kawecki 1994, Stearns et al. 1995). In the literature, the steep response to environmental variation is considered to be the presence of phenotypic plasticity, while the flatter response is supposed to be a sign of trait canalization (Van Buskirk and Steiner 2009). In this study, we show a clear example of plasticity *vs*. canalization, for the case of body size response to temperature.

The non-uniform pattern of plastic body size response across the thermal range has been confirmed for a number of protist species and has been termed the ’optimal thermal range for TSR’ (Walczyńska et al. 2016), the existence of which was then also confirmed by Blanckenhorn et al. (2021) for terrestrial insects. The theoretical scheme for the plasticity of body size across the thermal range from stressfully low to optimal to stressfully high, derived directly from figure 4 in Walczyńska et al. (2016), is shown in Fig. 1. Basically, the idea is that both population growth rate and body size show typical and consistent patterns around T_min_ and T_opt_. In the case of T_min_, population growth rate is negative below and positive above, while body size is smaller below than above. In the case of T_opt_ both, population growth rate and body size are lower below and above than at this temperature. One of the aims of this study is to test the universality of the pattern of non-linear body size response to external temperature and its relationship to fitness.

**Fig. 1.**
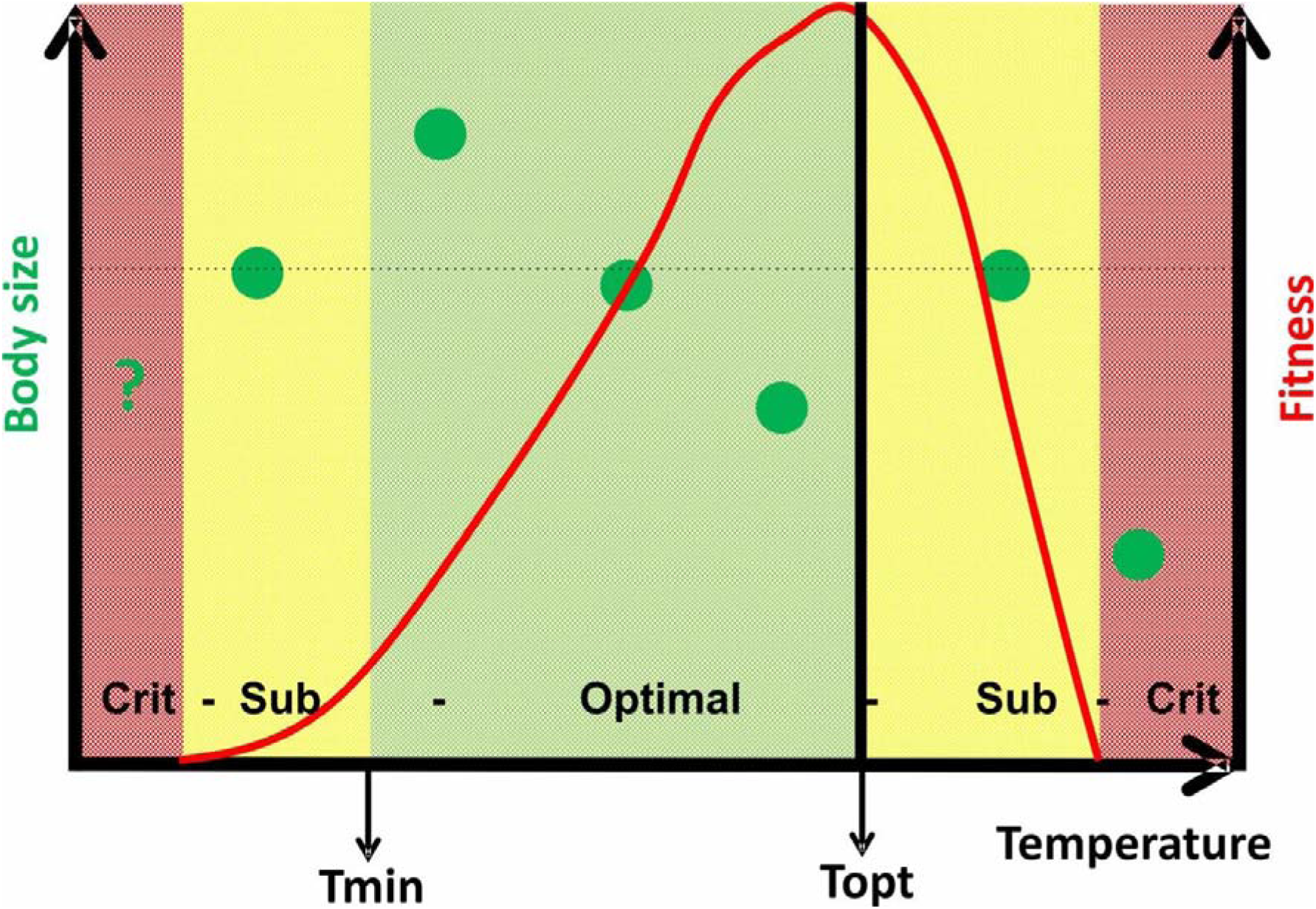
Conceptual background of the study, based on the previous results of Walczyńska et al. (2016). The plastic response of body size decrease with increasing temperature (green dots) is performed within a certain thermal range, which is limited between T_min_ and T_opt_, defined for fitness (red line). Beyond this range, under suboptimal conditions (“Sub”), body size is canalized (zero plasticity; dotted horizontal line). At stressfully high temperature (“Crit”), body size decreases even more, either to counteract negative physiological effects or as a result of such negative effects. The pattern for stressfully low temperature is unknown.

Our study of the conditions of the TSR performance involved different experimental populations of the rotifer *Lecane inermis* in order to relate the variability in TSR strength to organisms with different thermal preferences. *L. inermis* has previously been intensively studied with regard to the proximate and ultimate factors behind the TSR. Most importantly, the results showed that the TSR is an adaptation to temperature-dependent oxygen availability, because small rotifers had larger fitness (fecundity) than large rotifers under combination of high temperature and hypoxia (Walczyńska et al. 2015a). Therefore, it was justified to refer to TSR as an adaptation throughout this study. Furthermore, it was found that this growth pattern is performed both in the laboratory and in the field (Kiełbasa et al. 2014) and is both developmental and transgenerational (sensu McKenzie et al. 2021), as it is performed by the mother and during egg development (Walczyńska et al. 2015b). Recently, the TSR in *L. inermis* was found to be sensitive to environmental cues and under selective pressure (Walczyńska and Sobczyk 2022). Here we used five clones of *L. inermis* rotifer and three multiclonal experimental populations of *L. inermis*, the latter having previously been subjected to six months of experimental evolution with body size selection under different oxygenic conditions.

Our goals were: (i) to test for a common non-linear match between body size and fitness responses to temperature in experimental populations differing in their thermal history (Fig. 1), and (ii) to look for differences in the optimal thermal range for TSR between the studied populations and relate them to their thermal tolerance. Thus, this study aimed to contribute to the understanding of the mechanisms and limits of phenotypic plasticity. In particular, we address the issue of the complex pattern of body size response to temperature, and the sources of variability in the strength of this response.

## 2. Methods

### 2.1. Studied organisms

*Lecane inermis* Bryce (Rotifera, Monogononta) is a bacterivorous species. In the activated sludge ecosystem which it abundantly inhabits, this species loses its ability to reproduce sexually (Pajdak-Stós et al., 2014). Under standard conditions, the female lives for about 9-10 days, and the first offspring appear on the second or the third day after hatching (Miller 1931). *L. inermis* is able to proliferate at temperatures up to 38 °C (Walczyńska et al., 2016), and the generation time was previously estimated to be 2 days at temperatures between 15 °C and 25 °C (Walczyńska et al., 2015b).

We used eight experimental cultures of *L. inermis*. Four clones were among six previously examined by Stuczyńska et al. (2021). Additionally, we used a clone which has not been studied before in the context of the link between thermal performance curves and the strength of TSR response. These five clonal experimental populations are hereafter referred to as clonal groups (CG). The details on the results of the previous studies are provided in Table 1A. The other three experimental populations studied were the rotifers which had previously experienced the six-month experimental evolution in three different oxygenic conditions, hereafter referred to as post-evolution groups (PEG). PEGs were exposed to selection in conditions of constant normoxia (TN), constant hypoxia (TH), or temperature-dependent hypoxia (hypoxia only at high temperature) (TNH), under conditions of temperature fluctuating between 22 °C and 28 °C (Walczyńska and Sobczyk 2022). These groups are not clonal, because the experimental evolution was initiated with an even mixture of eleven clones of *L. inermis*. Details on PEGs are provided in Table 1B.

**Table 1.**
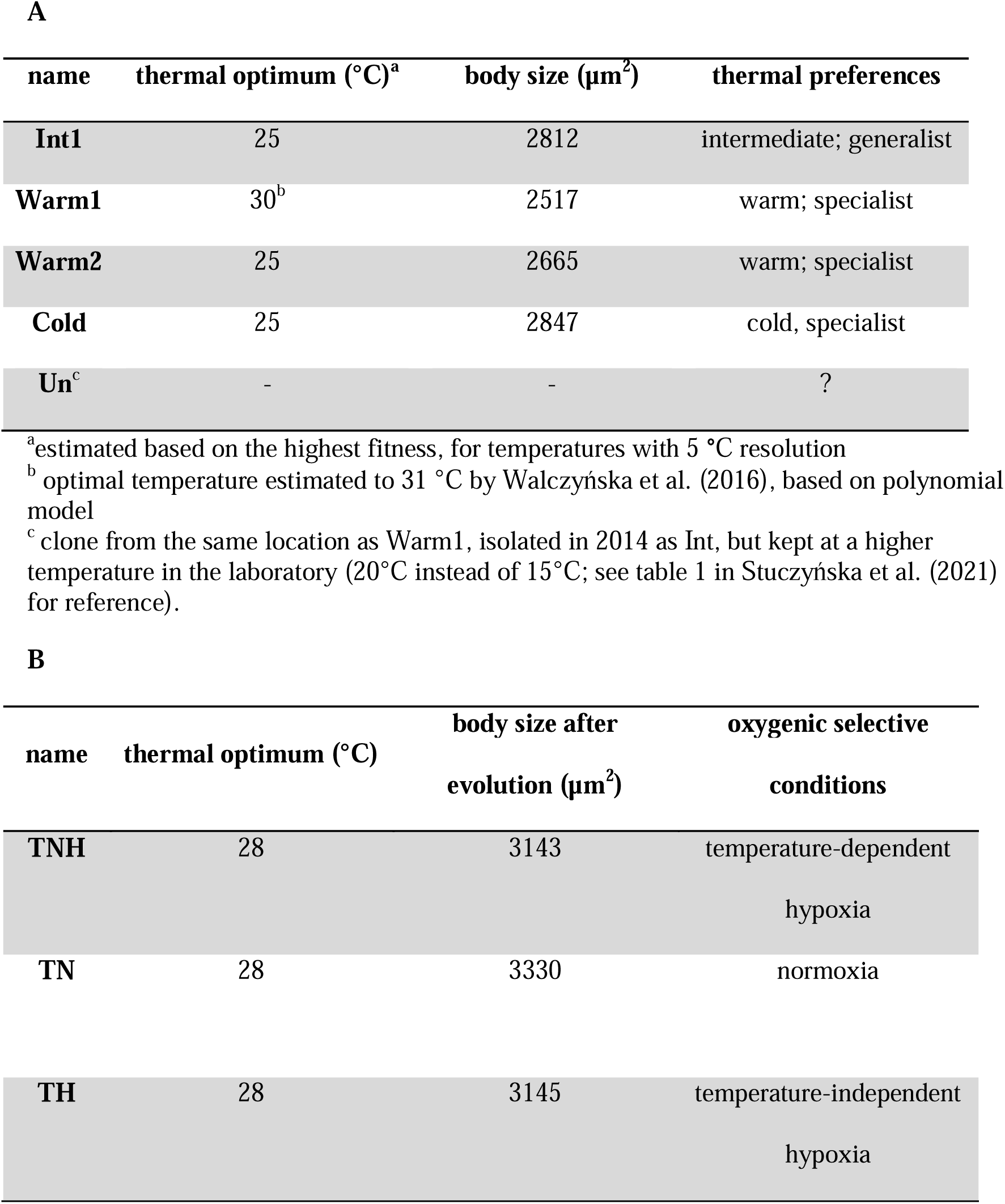
Description of experimental populations of the rotifer *Lecane inermis* used in this study. A – clonal groups (CG), data from Stuczyńska et al. (2021), no previous data for the Un clone; B – post-evolution groups (PEG), data from Walczyńska and Sobczyk (2022); thermal optimum – experimental temperature at which rotifers achieved highest fitness measured as the population growth rate; body size measured at standard temperature of 25 °C.

| name | thermal optimum (°C)<sup>a</sup> | body size (µm<sup>2</sup>) | thermal preferences |
| --- | --- | --- | --- |
| **Int1** | 25 | 2812 | intermediate; generalist |
| **Warm1** | 30<sup>b</sup> | 2517 | warm; specialist |
| **Warm2** | 25 | 2665 | warm; specialist |
| **Cold** | 25 | 2847 | cold, specialist |
| **Un<sup>c</sup>** | - | - | ? |

<sup>a</sup>estimated based on the highest fitness, for temperatures with 5 °C resolution
<sup>b</sup> optimal temperature estimated to 31 °C by Walczyńska et al. (2016), based on polynomial model
<sup>c</sup> clone from the same location as Warm1, isolated in 2014 as Int, but kept at a higher temperature in the laboratory (20°C instead of 15°C; see table 1 in Stuczyńska et al. (2021) for reference).

**B**
| name | thermal optimum (°C) | body size after evolution (µm<sup>2</sup>) | oxygenic selective conditions |
| --- | --- | --- | --- |
| **TNH** | 28 | 3143 | temperature-dependent hypoxia |
| **TN** | 28 | 3330 | normoxia |
| **TH** | 28 | 3145 | temperature-independent hypoxia |

In stock, all eight experimental populations were maintained in the 60 mm Ill Petri dishes in climatic chamber at 25 °C with Żywiec Zdrój® spring water (Poland) as a medium, fed with a nutrition powder used for rotifer mass culture NOVO® (patented by Pajdak-Stós et al. 2017), and a commercial bioproduct Bio-Trakt® (Zielone oczyszczalnie, Poland), a probiotic solution with known bacterial species composition, which created a stable microbial environment.

### 2.2. Study design

The experiment was divided into two stages. For logistical reasons, the study of CGs and PEGs was separated in time in the two stages. In the first stage (hereafter stage I), we tested for differences between the experimental populations in the optimal thermal range and in the strength of the TSR response over a wide temperature range represented by six treatments from 10 °C to 35 °C (Fig. 2). The body size results from this stage were used to analyse the strength of the TSR, while the population growth rate results were used to estimate the thermal performance curve. The combination of both traits was used to select the temperatures for the second stage of the experiment. In the next stage (hereafter stage II), we aimed to refine the estimates of optimal thermal ranges for TSR. Temperature thresholds within which the rotifers exhibited the clearest pattern of TSR were tested in a narrower resolution, at temperatures selected after stage I, based on the inverse pattern of body size around the low temperature threshold (Fig. 1). If body size was smaller at the lowest temperature than at the next-to-lowest temperature, we interpreted the result as T_min_ being between these two temperatures, and we used this lowest temperature and a temperature two degrees higher in stage II. If body size was greater at the lowest temperature than at the next-to-lowest temperature in stage I, we interpreted this as T_min_ being below this range. We took this lowest temperature and a temperature two degrees lower in stage II. We treated T_opt_ as the temperature at which the population growth rate, a direct measure of fitness, was maximal. We aimed to use three temperatures around T_opt_ to find the exact temperature in stage II. The result of this stage were used to estimate the thermal performance curves and to select the precise optimal thermal ranges for the TSR for each experimental population.

**Fig. 2.**
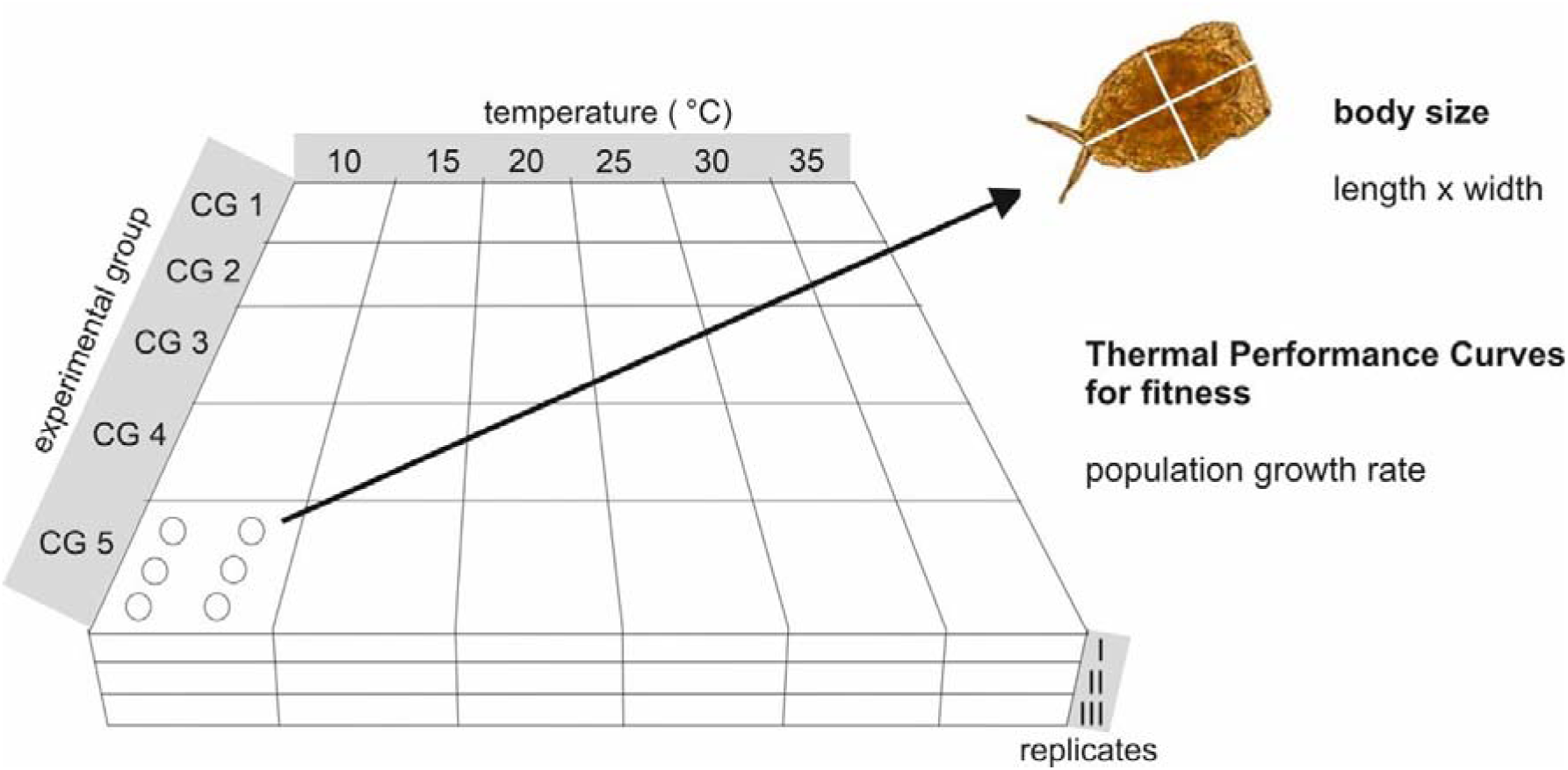
Experimental design for stage I. Each cell represents a combination of experimental population, temperature and replicate. For each cell, two rotifer traits were estimated: body size, calculated as length × width, and population growth rate, which was further used to model the thermal performance curve for fitness. Shown for clonal groups (CG), with post-evolutionary groups (PEG) having an identical design.

To initiate each experimental stage, rotifers were transferred to six-well tissue culture plates, 20±5 individuals per well, with 6 ml of Żywiec Zdrój® medium and 30 µl of Bio-Trakt® as food source. In stage II, rotifers were additionally fed with NOVO to enhance their proliferation. There were three replicates (= three plates) per combination of experimental population and temperature, with six wells within a plate acting as pseudoreplicates and analysed as pooled across wells (Fig. 2). This approach allowed the heterogeneity of conditions between wells to be considered.

### 2.3. Fitness and body size characterization

We examined two life history traits: (i) body size, which provided information on the plastic response to different temperatures (TSR), and (ii) population growth rate, which acted as a direct fitness measure and allowed the determination of the limits of the ’optimal thermal range for TSR’. For the first 2-7 days, depending on temperature, the rotifers were acclimated to new thermal conditions, to achieve both, rotifer proliferation and the stable age structure to secure the proper population growth rate estimation (Caswell 2001). For the population growth rate estimation rotifers were counted twice, after 2-9 days apart (for the highest and the lowest temperatures, respectively). The differences in the time gap allowed to account for the biophysical processes dependent on temperature. For the counting, 300 μl of culture was sampled from each well within a plate and transferred to an Eppendorf tube. Each well was refilled with 300μl of Żywiec Zdrój® water and fed with 30 μl of Bio-Trakt® (or 10 μl of Bio-Trakt® and *c*. 2 mg of NOVO in stage II). Therefore, one Eppendorf tube was prepared for each experimental population × temperature × replicate combination. The 25 μl subsamples were taken from each tube to estimate the rotifer density per 1 mL, following the standard method (Dubber and Gray 2009). Five samples were taken from each tube, the rotifers were counted using the light microscopy with different microscopes (3-4 counting people were involved at a time). The final rotifer number in a replicate was calculated from the three values remaining after removal of the highest and the lowest value. For the case of the lowest temperatures, the rotifers were counted directly from the wells (all rotifers per well). In such cases, the final rotifer number in a replicate (plate) was calculated as a sum of rotifers in all wells. To allow comparison of the density between the two counting methods, all densities were recalculated to 1 mL.

The population growth rate *r* was estimated as a difference between the number of rotifers from the second and the first counting, divided by the period of time between countings in days:

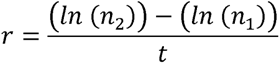

where: *r* – population growth rate, n_1_ – number of rotifers in first counting, n_2_ – number of rotifers in second counting, t – time period between two counting [days].

After the second counting, rotifers were fed. A day after, the samples were fixed with Lugol solution for body size measurements. Preserved rotifers, *c*. 30 individuals per replicate, were photographed using Olympus inverted microscope, Nikon DS-U1 camera and NIS Elements software with 40× magnification. Length and width of the rotifer lorica were measured in ImageJ (NIH, USA) and their multiplication acted as a proxy of area [μm^2^] (Fig. 2). This method was used previously in the studies on the TSR in *L. inermis* rotifers (Walczyńska et al. 2015a, Walczyńska et al. 2015b, Walczyńska et al. 2016, Walczyńska et al. 2017, Stuczyńska et al. 2021, Walczyńska and Sobczyk 2022).

### 2.4. Data analysis

All analyses were conducted via R programming language (R Core Team 2022) in RStudio (Posit team 2023), and in packages: “rTPC” for thermal performance curve calculations (Padfield et al. 2021), “lme4” for linear regression calculations (Bates et al. 2015) and “emmeans” for post-hoc tests (Lenth 2023). A “performance” package (Lüdecke et al. 2021) was used to test whether the assumptions for ANOVA and the linear models were met.

#### 2.4.1. Body size and strength of the TSR

A statistical model for testing the differences in body size from stage I included experimental population and temperature as categorical, fixed factors and replicate, nested in (group × temperature), as a random factor. To analyse the strength of the TSR, the linear mixed model was performed separately for each CG and PEG population for body size from stage I. The test was performed for the thermal range specific to each experimental population. To secure the range within which body size decrease is assumed to be linear, the range was enclosed between the lowest temperature at which the mean population growth rate *r* was positive and the temperature no higher than that at which the population growth rate was maximal in stage I. The slope of the decrease in body size with increasing temperature was treated as equivalent to the strength of the TSR response. The relationship between the strength of the TSR and thermal preferences was tested for CGs (no sufficient sample size in PEGs) in simple regressions relating slope of body size response over temperature to thermal range and to T_opt_, separately.

#### 2.4.2. Thermal performance curve and optimal thermal range

TPC was estimated using two methods. The first estimation was based on tracking the body size pattern, as presented in Fig. 1. The inverse body size pattern informed about T_min_ and T_opt_. The second estimation was based on formal model fitting. In this case, we tested all the models for TPCs estimation suggested by Padfield et al. (2021) and applied those that fit the data reasonably well based on visual assessment, and were common to all experimental populations: Briere2_1999, Joehnk_2008, Kamykowski_1985 and Lactin2_1995. The parameters estimated for each model separately are displayed in supplementary Table S1. The final estimates of the T_min_ and T_opt_ parameters were calculated as averages across these models. In both methods applied, the range of thermal tolerance was calculated as a difference between T_opt_ and T_min_.

Finally, the existence of a canalization of body size beyond the ‘optimal thermal range’ was tested. In stage I, when the reverse body size response was observed around T_min_ (Fig. 1), the difference between body size at the lowest temperature was compared with that at temperatures above T_opt_ using ANOVA with temperature as a categorical, fixed factor.

## 3. Results

### 3.1. Descriptive results

The mean body size for each experimental population over temperature achieved in stage I is presented in Table 2. The relative differences in size between CGs were consistent for all except clone Int, which achieved a relatively smaller size in the present study compared to the previous one (bolded column in Table 2 *vs*. Table 1). As predicted and found in the previous study, clone Warm1 was the smallest and clone Cold the largest. The new clone Un had a body size intermediate between warm-preferring clones and cold-preferring clone. The relative differences in body size between PEGs were not consistent with the previous results (Table 2 vs. Table 1); comparing with the previous results, TN and TNH were now similar in size while TH was smaller.

**Table 2.**
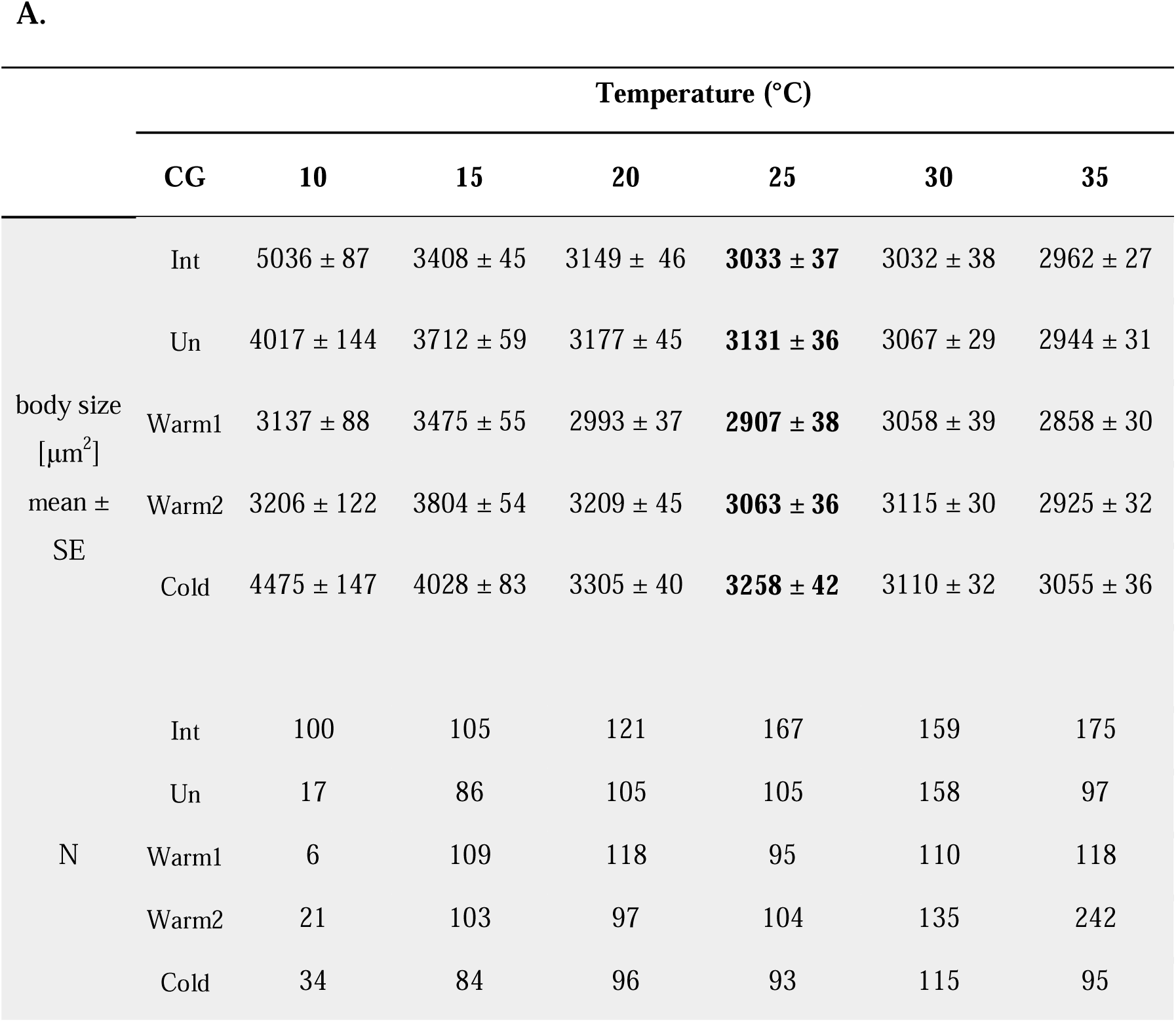
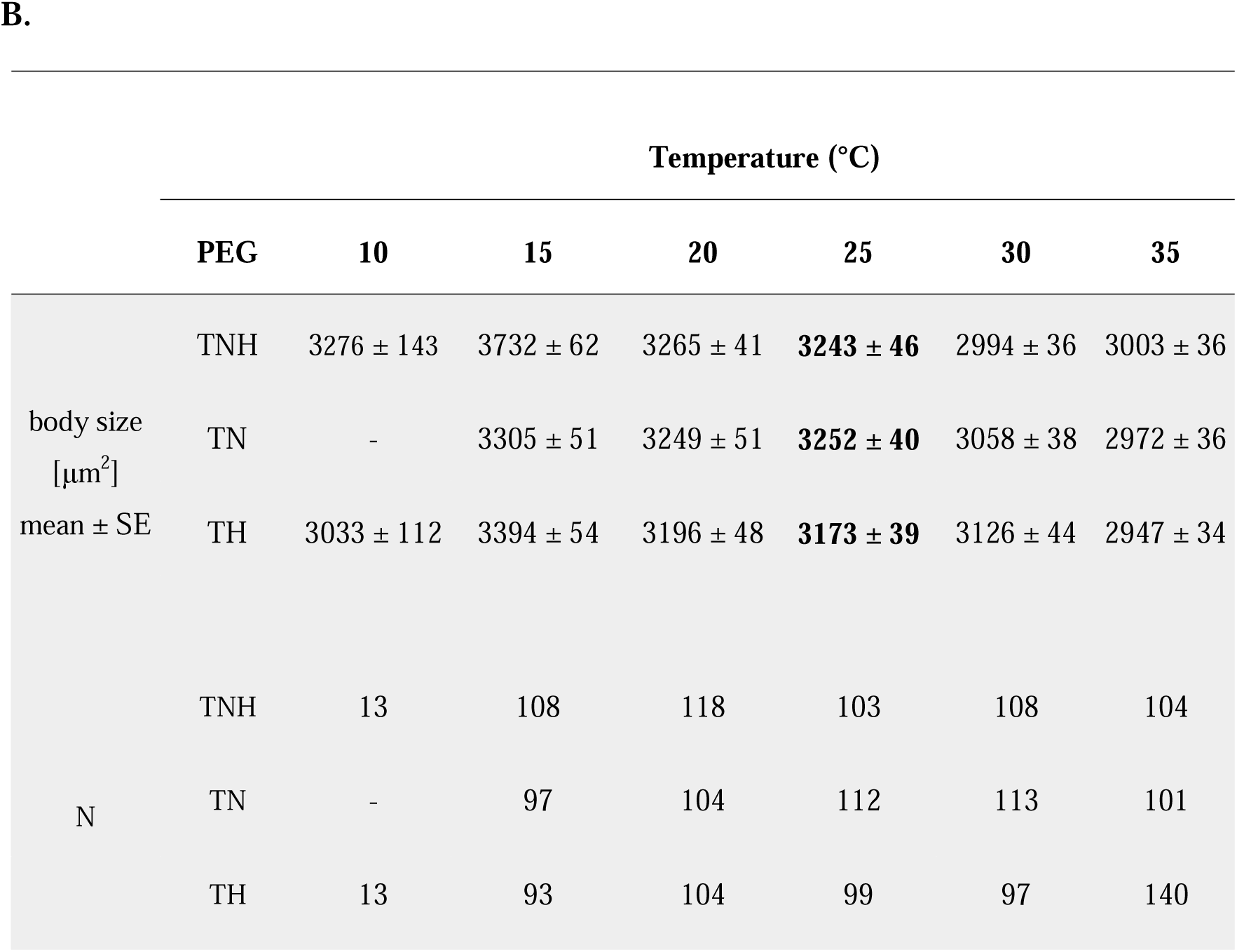
Body size for each CG (A) and PEG (B) achieved in experimental stage I over six temperatures. The bolded temperature is for comparison with the previous data in Table 1. N – sample size

The population growth rate *r* achieved in experimental stages I and II is shown in supplementary Fig. S1. The results of stage II were comparable to those of the previous studies (Stuczyńska et al. 2021, Walczyńska and Sobczyk 2022). Stage I rotifers did not grow well at 25 °C and 30° C, and partly at 20 °C, and the most likely reason for this was microbial infection. However, the pattern of lower than expected numbers of individuals was observed in the second counting and not in the first, and no dead rotifers were observed during the counting, meaning that infection may have occurred just before the second counting and may have affected the egg laying rather than the plastic response of body size to temperature. The population growth rate calculated for stage I was not comparable to that achieved in stage II because infection may have caused relatively lower proliferation at intermediate temperatures in stage I and because NOVO was added to the diet in stage II. We decided to estimate TPCs for population growth rates pooled for stages I and II, but without *r* values for 20 °C, 25 °C and 30 °C in stage I (Fig. 3). All mean population growth rate data are presented in Fig. S1.

**Fig. 3.**
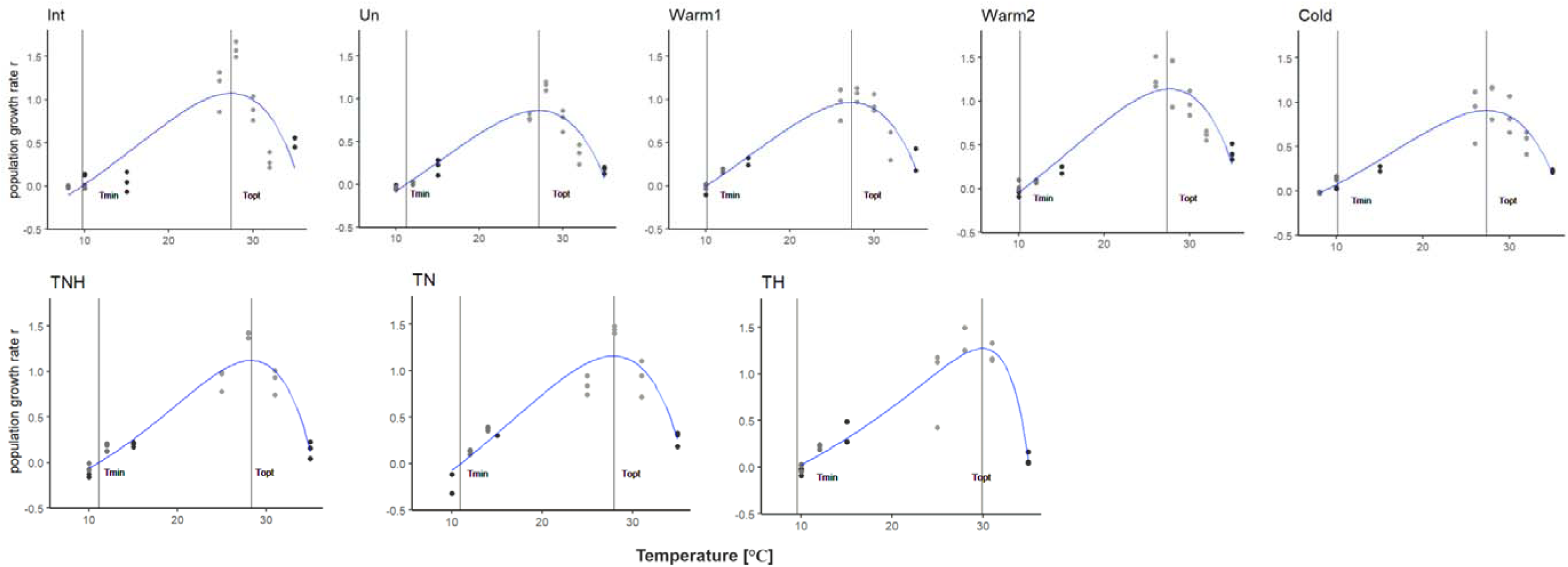
Population growth rate data for all experimental populations achieved in stage I (black dots) and stage II (grey dots), excluding data for temperatures 20 °C, 25 °C and 30 °C in stage I. The thermal performance curve (blue line) and the corresponding T_min_ and T_opt_ are estimated using the Lactin2_1995 model (Padfield et al. 2021).

### 3.2. Body size pattern in stage I and selecting temperatures for stage II

In both CGs and PEGs, body size differed significantly between experimental populations and these differences varied across temperature, as shown in significant interaction of temperature × exp. population (CGs: F_(4, 3044)_ = 21.15, p < 0.001; PEGs: F_(2, 1620)_ = 7.75, p < 0.001). The effect of replicate was significant for CGs (p = 0.023) and not significant for PEGs (p = 0.162). In stage I, the reverse pattern of body size response to temperature was observed for the two lowest temperatures of the clones Warm1 and Warm2 (Fig. 4A). The interpretation of such a result is that for these two clones, T_min_ is between 10 °C and 15 °C, while for all other clones, T_min_ is below 10 °C. However, as the population growth rate at 10 °C was negative for Warm1, Warm2 and Un (Fig. 3), the temperatures used to select the more accurate T_min_ in stage II were chosen to be 10 °C and 12 °C for these three clones and 8 °C and 10 °C for Int and Cold. In the PEGs, all three populations showed a pattern indicating a T_min_ between 10 °C and 15 °C (Fig. 4B). In the case of TN, the rotifers did not proliferate at all at 10 °C. Therefore, in stage II, 10 °C and 12°C were used to determine the exact T_min_ for TNH and TH, and 12 °C and 14 °C were used for TN.

**Fig. 4.**
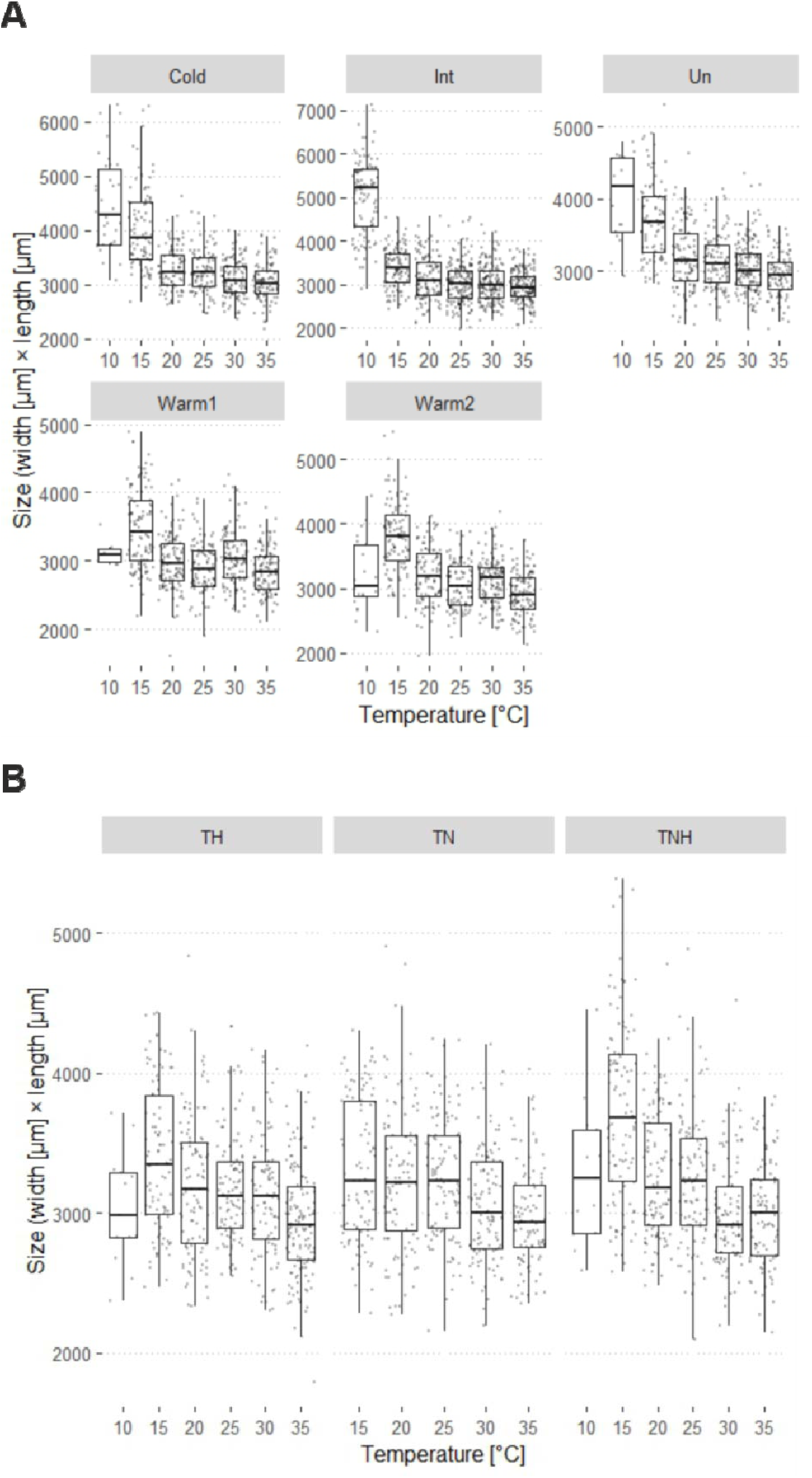
Boxplots for body size of rotifers exposed to six temperatures in stage I, with individual data points jittered. Median with lower and upper quartile and extremes as whiskers. Dots represent jittered measurements of individual body size. A – CGs, B – PEGs

For T_opt_, the patterns were more complex and both CGs and PEGs showed comparable body size at highest temperatures (Fig. 4), while referring this pattern to *r* was not possible because of the possible microbial infection affecting the population growth rate (Fig. S1). Therefore, based on the previous data on T_opt_ in PEGs (Table 1), we used three temperatures around 28°C (26 °C, 28 °C and 31 °C) in stage II. For CGs we decided that using four temperatures around T_opt_ would be more effective in selecting the exact value and in stage II we used 26°C, 28 °C, 30 °C and 32 °C.

### 3.3. Body size in stage II and estimation of T_min_ and T_opt_

Following the reasoning from Fig. 1, the significantly smaller rotifers were found at lower than at higher temperatures in the T_min_ estimation in all experimental populations except Cold (significantly larger rotifers at lower temperature), and Warm1 (no difference in body size between two temperatures; Table S2 and Fig. 5). The pattern of *r* is largely consistent with predictions and growth was slower below than above T_min_. Exception was clone Int, where the two values were comparable. We interpret these results to mean that in the case of Cold, T_min_ is lower than that set in stage II, whereas in the case of Warm1, the narrowed temperature range of stage II must not have been well set compared to the wider range of stage I. For those experimental populations which showed the increasing body size pattern around T_min_, we assumed T_min_ to be the value between the two temperatures set in stage II. For the Cold clone, we assumed T_min_ to be 8 °C (population growth rate was around 0 for this temperature; Fig. 5), and for the Warm1, we assumed T_min_ to be the mean of the thermal range in stage I, that is, 12.5 °C. All T_min_ values are shown in Table 3.

**Fig. 5.**
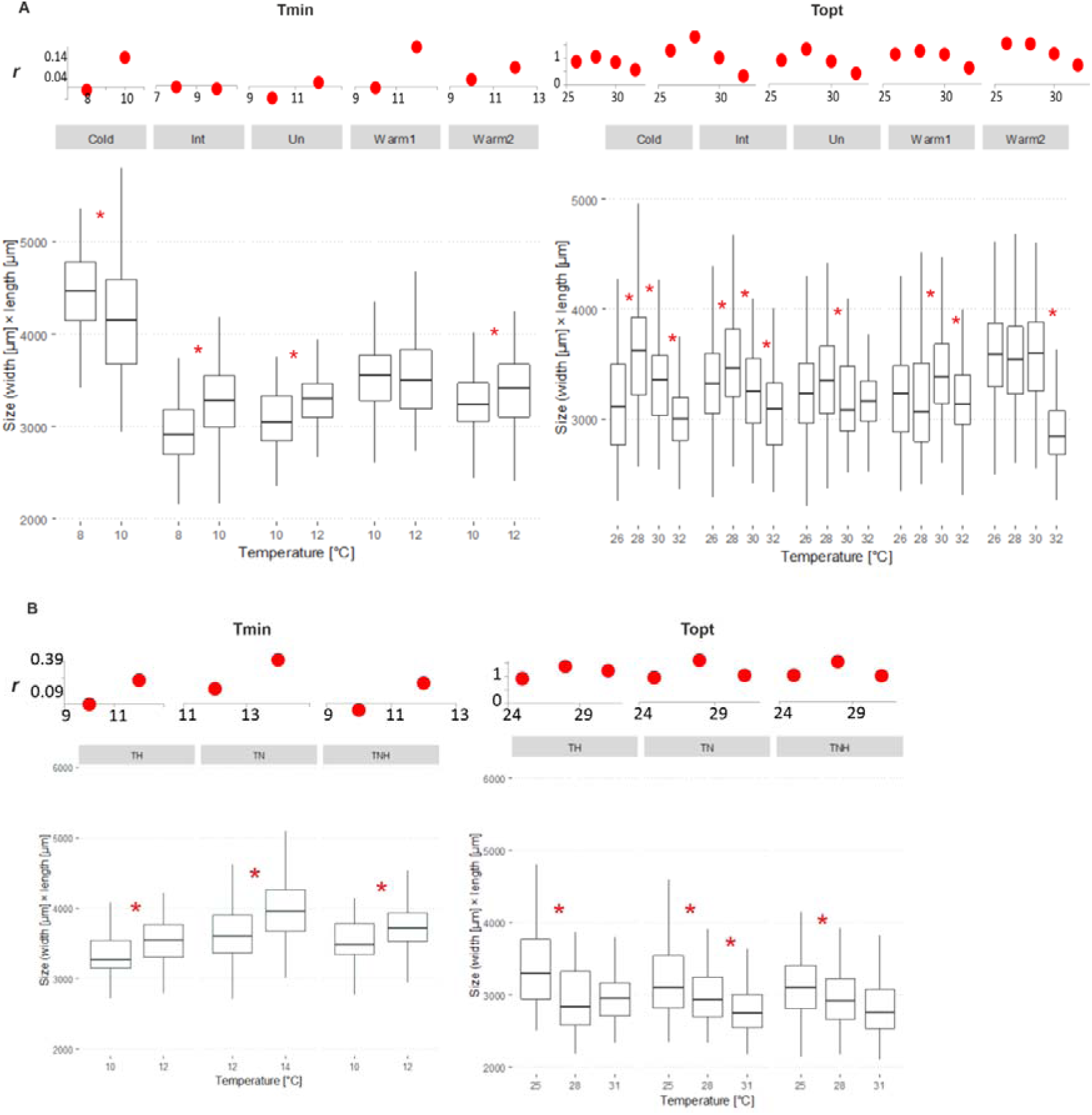
Population growth rate *r* (upper figures, red dots) and body size (lower figures, boxplots) for temperatures used in stage II to delineate T_min_ (left) and T_opt_ (right). Asterisks denote a significant difference between the two neighbouring temperatures. Median with lower and upper quartile and extremes as whiskers. A – CGs, B – PEGs

**Table 3.** Parameters describing the TPC for population growth rate, given as raw data (r_max_) and estimated from the body size pattern according to Fig. 1 (a), or formally estimated as averages from four mathematical models (b). r_max_ – maximum population growth rate, T_min_ – temperature around which population growth rate reaches zero, T_opt_ – temperature at which population growth rate is the highest. CG – clonal groups, PEG – post-evolution groups

| | $r_{\max}$ | | $T_{\text{opt}}[^{\circ}\text{C}]$ | | $T_{\min}[^{\circ}\text{C}]$ | | thermal range | |
| --- | --- | --- | --- | --- | --- | --- | --- | --- |
|  | a | b | a | b | a | b | a | b |
| **CG** |  |  |  |  |  |  |  |  |
| Int | 1.58 | 1.08 | 27 | 26.9 | 9 | 9.8 | 18 | 17 |
| Un | 1.16 | 0.88 | 27 | 26.5 | 11 | 11.3 | 16 | 15 |
| Warm1 | 1.06 | 0.98 | 29 | 26.5 | 12.5 | 10.3 | 16.5 | 16 |
| Warm2 | 1.30 | 1.15 | 29 | 26.8 | 11 | 10.6 | 17 | 16 |
| Cold | 1.04 | 0.91 | 27 | 27.1 | 8 | 8.6 | 19 | 18 |
| **PEG** |  |  |  |  |  |  |  |  |
| TNH | 1.40 | 1.11 | 28 | 28.3 | 11 | 11.1 | 17 | 17 |
| TN | 1.45 | 1.15 | 28 | 27.9 | 13 | 11.0 | 15 | 17 |
| TH | 1.38 | 1.26 | 28 | 30.1 | 11 | 9.6 | 17 | 20 |

As for T_opt_, the body size patterns in the CGs showed a significant increase between 26 °C and 28 °C for clones Int and Cold and a tendency for clone Un with p = 0.0835 (Table S2), or between 28 °C and 30 °C (Warm1), with no increase in the case of Warm2. The significant body size decrease was observed above 28 °C (Int, Un, Cold), or above 30 °C (Warm1, Warm2; Fig. 5). The referential *r* pattern confirmed the concave shape for the thermal range examined in all experimental populations, and the decline in growth rate at the highest temperatures was particularly evident (Fig. 5). For Int, Un, Cold, and Warm1 the final T_opt_ was assumed to be the mean between two temperatures at which an increase in body size was observed. In the case of Warm2, where no such increase was observed, the T_opt_ was assumed to be the same as for Warm1, as the similar pattern of body size decrease at higher temperatures (= tolerance to higher suboptimal temperature) was shared by these two clones. For all PEGs, body size decreased above 25 °C (Fig. 5). According to Fig. 1, such a result would imply that T_opt_ is closer to 25 °C than to 28 °C. However, this result contradicted the fitness data, where all PEGs performed best around 28 °C, which was consistent with the previous study (Walczyńska and Sobczyk 2022). Therefore, in PEGs, T_opt_ was assumed to be 28 °C and we discuss this decision below. All T_opt_ values and additional values for the highest *r* values, r_max_, and the thermal ranges as the difference between T_opt_ and T_min_, are shown in Table 3.

The results for the TPCs formally estimated on the basis of four models showed only slightly different values for all three measures, r_max_, T_min_ and T_opt_, compared to the method based on tracking body size changes (Table 3). The largest discrepancies in the comparison of the two methods concern T_opt_. The relative differences deviated for Warm1 and Warm2, where T_opt_ was considerably higher for the method of body size tracking, and for TH, where T_opt_ was lower for this estimation method. This pattern is discussed below.

We showed that body size was smaller below T_min_ than above, which was consistent for almost all experimental populations (Fig. 5). For T_opt_, the results were more complex. For CGs, the results obtained using the body size tracking method were consistent with the conceptual predictions from Fig. 1 and with previous results (Stuczyńska et al. 2021). The results of the formal analysis were similar for all CGs, except for the two Warm clones, where the analysis showed a lower estimate. For PEGs, the pattern of body size did not match that predicted in Fig. 1; body size decreased across the three temperatures studied. Based on the visual representation (the vertical line denoting T_opt_ in Fig. 3), the formal estimation of T_opt_ was accurate for TN and TNH and gave an inaccurate (too high) value for TH.

### 3.4. Strength of the TSR and canalization of body size

In all experimental populations, body size decreased significantly with increasing temperature, within the individually set ’optimal thermal range’ (Table 4). The slopes obtained for the CGs were very similar to those previously reported (Table 2 in Stuczyńska et al. 2021). In absolute terms, the Int clone showed the steepest slope as expected, while all three specialised clones (Cold, Warm1 and Warm2) showed comparable, flatter responses. The Un clone showed the flattest response of all CGs. PEGs showed flatter slopes than CGs. The slopes for TSR calculated at 22 °C, 25 °C and 28 °C, for the data shown in figure 4 in Walczyńska and Sobczyk (2022), are -0.028, -0.018 and -0.019 for TNH, TN and TH, respectively. Although the slopes of response obtained in this study were much flatter, the relative differences between them and the previous ones were maintained (Table 4). Simple regression analysis for CGs showed neither the relationship of slope with thermal range (t = - 1.26, R^2^ = 0,13, p = 0.30) nor with T_opt_ (t = 0.64, R^2^ = -0.18, p = 0.57; Fig. 6).

**Fig. 6.**
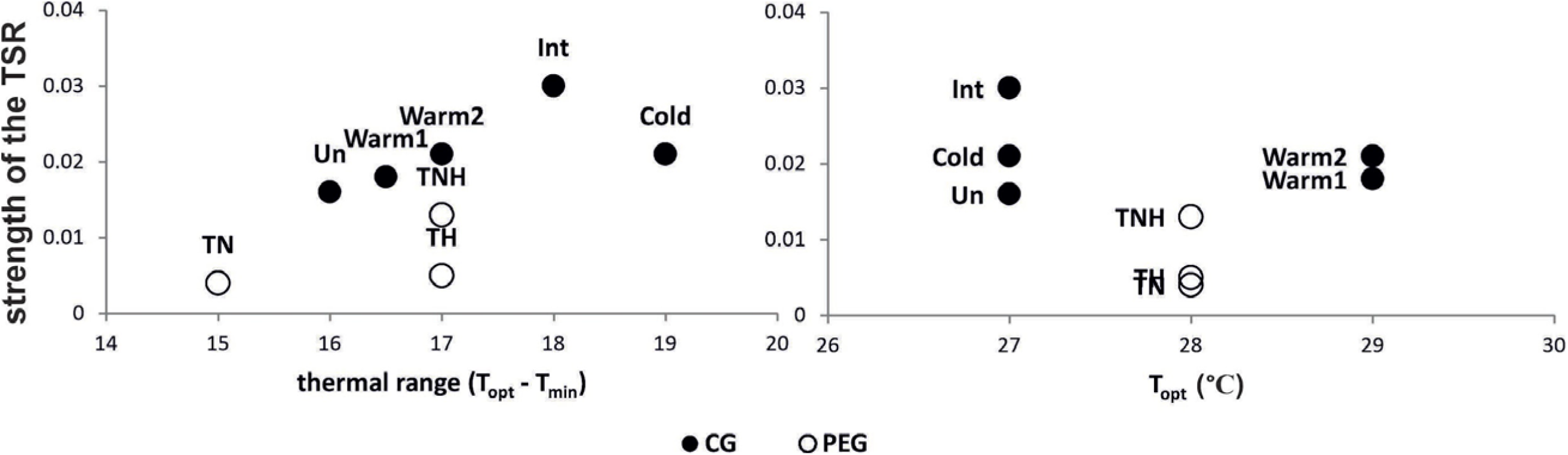
The relationship between the strength of the TSR and two measures of thermal preference: range of temperature tolerance (left) and T_opt_ (right) informing about the thermal specialisation. For clarity, the strength of the TSR is shown as the modulus of the negative slope, [slope]. CG – clonal experimental populations, PEG – post-evolution experimental populations; the acronyms of the experimental populations are defined in Table 1.

**Table 4.** Details of the response of body size to temperature (the temperature-size rule pattern), estimated for temperatures in the range between T_min_ and T_opt_, selected in the formal analysis for each experimental population separately. CG – clonal group, PEG – post-evolution group

| | slope | $R^2_{\text{adj}}$ | df | t | p |
| --- | --- | --- | --- | --- | --- |
| **CG** |  |  |  |  |  |
| Int | -0.030 | 0.47 | 491 | -20.8 | < 0.0001 |
| Un | -0.016 | 0.17 | 294 | -8.0 | < 0.0001 |
| Warm1 | -0.018 | 0.18 | 320 | -8.5 | < 0.0001 |
| Warm2 | -0.021 | 0.29 | 302 | -11.1 | < 0.0001 |
| Cold | -0.021 | 0.33 | 305 | -12.3 | < 0.0001 |
| **PEG** |  |  |  |  |  |
| TNH | -0.013 | 0.19 | 435 | -10.3 | < 0.0001 |
| TN | -0.004 | 0.03 | 424 | -3.5 | 0.0005 |
| TH | -0.005 | 0.03 | 391 | -3.7 | 0.0002 |

## 4. Discussion

The contribution of this study to the issue of operating conditions and variability in phenotypic plasticity of body size response to temperature is to (i) confirm the existence of clearly describable conditions under which this plastic response operates, a concept previously proposed (Walczyńska et al. 2016) and developed more extensively in this study, (ii) discuss possible reasons for the variability in the strength of the TSR response, and (iii) discuss the extent of evolutionary conservatism of thermal tolerance descriptors by studying closely related organisms with different thermal histories.

### 4.1. Correspondence between body size and fitness across temperature

The first objective of our study was to test whether the non-uniform pattern of body size response within and beyond the ‘optimal thermal range for TSR’, previously found by Walczyńska et al. (2016) and presented in Fig. 1, was consistent in different experimental populations, either clonal or multiclonal, representing different thermal histories. We showed the clear confirmation for the pattern of body size around T_min_ for population growth rate, consistent for almost all experimental populations (Fig. 5). Referring to T_opt_, the results for CGs generally followed the predicted pattern, especially those estimated using the body size tracking method, while PEGs did not. There are two possible sources of deviation of PEGs from the theoretical predictions of the body size tracking method. The first relates to the genetic composition of the PEGs, which consist of the unknown number and composition of different clones. The pattern may have differed from predictions due to competition within these clonal mixtures. However, the fitness pattern contradicts the body size pattern, as all three PEGs had the highest fitness around 28°C as predicted (Fig. 3). This makes the second source of the observed body size variation more likely, namely that the resolution of the temperatures selected in stage II was not sufficient to identify the exact T_opt_. For these reasons, we decided to select T_opt_ based on fitness rather than body size patterns. However, when considering T_min_ and T_opt_ in CGs and PEGs, it is worth noticing that T_opt_ in PEGs was the only case where body size and fitness did not show the expected pattern. It is worth noticing that less consistent comparison of the two estimation methods regarding T_opt_ than T_min_ could have resulted from the previously reported considerable variability of asymmetry in TPCs around T_opt_ (Buckley et al. 2022). In addition, considering the flatter shapes for fitness around T_opt_ for specialised clones than for a generalist (Int), we suggest that specialised organisms have a wider thermal range around the temperature at which they perform best than generalists. This idea requires further investigation.

Canalization, i.e. the lack of plastic response, of body size outside the ’optimal thermal range’ could only be analysed for two CGs (Warm1 and Warm2) and two PEGs (TH and TNH), which showed a clear pattern of smaller body size below than above T_min_ in stage I. The analyses confirm the schematic assumption from Fig. 1. In all but TNH, there was no difference in body size between the lowest and next highest temperature, which was above the optimum, while at the highest temperature body size value was significantly lower than that at the two lower temperatures, meaning that 35 °C was stressful for the rotifers studied. In TNH there was no difference in body size between 30 °C and 35 °C. However, it is possible that the stressful effect of too much heat is more pronounced at lower temperatures (closer to 30 °C) in this experimental population than in the other two. For the specific case of phenotypic plasticity examined in the former (Walczyńska et al. 2016) and in this study, we have shown that the slope of the response may differ between organisms, reflecting their responsiveness, whereas canalization is a separate, clear and unambiguous phenomenon of zero plasticity.

Therefore, in this study we show an example of environmentally-driven, conditional induction of canalization. Such conditional changes between plasticity and canalization are evolutionary justified as both are costly under different ecological contexts (Van Buskirk and Steiner 2009). Phenotypic variation may be canalized by physiological mechanisms that are responsible for maintaining the physiological homeostasis, as suggested by Oleksiak and Crawford (2012). Evidence of metabolic canalization in response to environmental change, confirming such a mechanism, has been found in fish (Norin et al. 2016). With regard to the canalization of body size under suboptimal conditions, a mechanism based on cell size adjustment has been proposed by Walczyńska et al. (2018).

### 4.2. Variation in the strength of the TSR response

The second objective of this study was to relate the strength of the TSR to the thermal tolerance characteristics of experimental populations. A study of interclonal variation in life-history traits in *Daphnia magna* showed that maternal, developmental and environmental factors were the three main players (Harney et al. 2017). All of these sources of variance have been previously reported for TSR performance in rotifers (Walczyńska et al. 2015b, Walczyńska and Sobczyk 2022). Our novel, previously suggested contribution is that the strength of TSR might be related to the generalist-specialist strategy continuum (Walczyńska and Serra 2014a). This relationship was confirmed for six clones of the *L. inermis* rotifer (Stuczyńska et al. 2021). Separate experiments for CGs and PEGs prevented a joint analysis, and the number of experimental populations in CGs only was sufficient to conduct a formal analysis of the slope of the TSR response and thermal range or T_opt_. We did not find significant effects, but the graphical presentation of the results (Fig. 6) shows that it is not justified to reject the existence of such relationships (Fidler et al. 2006). The small sample size has a negative impact on the power of the analysis and any outlying point has a considerable effect on the estimate. In the case of the relationship between the strength of the TSR and the thermal range of the CGs, such a deviating point is for the Cold clone. In the case of T_opt_, the main problem is a low accuracy of estimation. The general pattern, which can only be based on trends shows that (i) the TSR response is stronger in clonal (CGs) than in multiclonal experimental populations (PEGs), and (ii) there is a tendency for a stronger TSR response in clonal experimental populations characterised by a wider range of thermal tolerance.

The above trends, together with the previous results (Walczyńska and Serra 2014a, Stuczyńska et al. 2021), indicate that it is worth further exploring whether organisms with different levels of thermal specialisation differ systematically in their plastic response to temperature. Such studies are needed in order to reach a final conclusion about the differential vulnerability of generalists or specialists to global warming (Kingsolver 2009, Huey et al. 2012), especially in light of recent evidence on the distinct and complementary roles that specialists and generalists play in natural communities (Dehling et al. 2021). Our results suggest that generalists may have a stronger plastic response to temperature change, which may make them less sensitive to abrupt temperature changes than specialists. On the other hand, the idea suggested above that specialists have a flatter area around T_opt_ than generalists suggests that the resolution of thermal change is equally important, as small changes around T_opt_ may be less harmful to specialists than to generalists.

### 4.3. Body size and thermal tolerance

Body size is considered to be the most important life-history trait because it is the result of other life-history traits, growth and development, that have been evolutionarily optimised against external mortality (Kozłowski 1992, 2006), and because it is the first-line trait on which natural selection acts (Hildrew et al. 2007). This importance is reflected in the fact that body size appeared to be an accurate predictor of species niche in biodiversity modelling (Scheffer et al. 2018). Body size has also been suggested to correspond to heat tolerance (Leiva et al. 2023). In fact, this correspondence is even broader for rotifers, as body size was previously found to be related to general thermal conditions in the living habitat (Walczyńska et al. 2021), and to thermal conditions at hatching from resting eggs (Walczyńska and Serra 2014b). More importantly for the present study, body size was found to be associated with thermal preferences at the clonal level (Stuczyńska et al. 2021). Comparing the data in Table 1, which refers to the previous studies, with Tables 2 and 3 from this study shows that we have confirmed this relationship. The same clones are the smallest and most warm-preferring here and there, while the largest clone again appeared to be the largest and most cold-preferring. This consistency was not confirmed for the multiclonal rotifer groups (PEGs) that had previously been exposed to body size selection under different oxygen conditions. However, it is important to note that the period between the completion of the experimental evolution and the start of this study (2.5 years) was much longer than the experimental evolution itself (six months), and during this period of relaxed selection the interclonal competition could have taken place. This may be also the reason why the PEGs showed weaker response (smaller slope modulus) in this study as compared to the previous one; the physiological plasticity in these clone mixtures could have been reduced in a stable environment (Morgan et al. 2022). This difference between CGs and PEGs may have important implications for understanding the effects of exposure to temperature change in populations with different genetic compositions.

The Un clone is an exception throughout our study as it is characterised by a relatively large body size, high T_min_, low T_opt_ (= narrow thermal tolerance range) and weak TSR response. This may be due to a specific physiological adaptation to a particular parameter related to the activated sludge environment from which the clone originated, or to some kind of handicap that allows this clone to propagate in the laboratory but would prevent it from proliferation in the field.

### 4.4. Repeatability of organismal response to temperature

Interestingly, we found a high degree of consistency in the relative differences in the traits investigated in the previous and current studies of the experimental populations. Apart from the comparison for body size described above, this repeatability also refers to T_min_ and the strength of the TSR response over an ‘optimal thermal range’. In terms of T_min_, the CGs assumed to be more warm-preferring according to the previous study still had the lowest tolerance to cold (= highest T_min_), followed by the Int clone and the most cold-tolerant Cold clone. PEGs, previously selected at a relatively high temperature, had a T_min_ similar to that of the warm-preferring CGs, as expected. In terms of the strength of the TSR response, not only did all the experimental populations show similar relative differences in the slope of the body size response to temperature as previously found, but the CGs also showed very similar slope values now and then (Stuczyńska et al. 2021). Considering how sensitive phenotypic plasticity responses are to both, different physiological states of organisms and external factors (nutritional, aerobic, infectious), the level of consistency in these results is intriguing.

The relatively high consistency of TPC descriptors across studies has been previously reported for *Drosophila melanogaster* (Kellermann et al. 2019), while Pallares et al. (2021) linked TPC patterns of the water beetle *Enochrus jesusarribasi* to the thermal history of the organism. Another measure of thermal tolerance, CT_max_, was also found to be associated with thermal history in *Drosophila melanogaster* (Kellermann et al. 2017) and in the fish *Galaxias zebratus* (Olsen et al. 2021). This is not surprising, given that all, acclimation, developmental plasticity and transgenerational plasticity, influence the shape of TPCs (Schulte et al. 2011). Our study empirically confirms the importance of organismal thermal history, in our case, related to the laboratory maintenance, in predicting the details of its thermal tolerance. In experimental clonal populations, thermal tolerance descriptors are relatively consistent even after long periods of maintenance under mild and stable laboratory conditions. Such a long-term effect may be explained by the consistency of individual metabolic rates, as previously found in the lizard *Lampropholis delicata* (Kar et al. 2022). The multi-clonal experimental populations show less consistent results, although, interestingly, the relative strength of the plastic response compared between two studies about two years apart was maintained. Further studies on different organisms would be needed to confirm whether this is an incidental finding or a more general rule. The general pattern of thermal adaptation resulting from the thermal history of organisms merits further study, as current results do not provide a clear answer to the nature and strength of their relationship (Aguado and Clusella-Trullas 2021, Santos et al. 2023).

### 4.5. Usefulness and limitations of thermal performance curves

In our study, two methods were compared in the delineation of T_min_ and T_opt_ for the fitness measure, (i) body size tracking and (ii) formal analysis of TPC shape estimation. Despite some inconsistencies, both methods showed relatively similar results, which is particularly noteworthy given that the formal estimation was performed on population growth data from two experimental stages with different feeding regimes. Such a result suggests that TPC estimation provides reliable information if only the sufficiently wide thermal range is considered for an organism with sufficiently well-known thermal preferences. The lesson from our study is that thermal resolution should be as low as possible in order to obtain accurate T_min_ and T_opt_.

There is a lively discussion about the usefulness and limitations of thermal performance curves as a tool for predicting the effects of global warming on ectotherms (e.g., Kellermann et al. 2019, Khelifa et al. 2019, Blanckenhorn et al. 2021, Pallares et al. 2021, Malusare et al. 2023). It seems that most of the problem is the complexity and context dependency of the pattern, which prevents the comparison of the shape of TPCs using the same mathematical models. Shi and Ge (2010) and more recently Padfield et al. (2021) have provided a comprehensive description of ready-to-use methods for estimating TPCs using different mathematical models. However, it is clear from the literature that there are differences between experiments in terms of the most appropriate model to be used. For example, Angilletta (2006) suggested the Gaussian function as the best to describe the TPC shape, Shi and Ge (2010) recommended Performance, Brière1 and Brière2 models for this purpose, while Pallares et al. (2021) used the Flinn_1991 model as the final one after the formal model selection. In our study, only the Brière2 model was found to be useful for our data from among those mentioned above. However, this model showed a very similar estimation to three other models (Table S1). These examples show that although we are much closer to being able to more accurately estimate the shape of the TPC for a given study, we are still far from being able to compare them between studies. Information on the sources of variation between TPCs, such as that provided in this study, may contribute to improvements in this regard.

## 5. Conclusions

Phenotypic plasticity is an elusive phenomenon, extremely sensitive to internal (physiological; e.g., Seebacher et al. 2015) and external factors, and therefore very difficult to study and, more importantly, to understand the conditions under which it operates (Via 1993). Especially important in this regard is a phenomenon of phenotypic plasticity in body size response to changing temperature, because decrease in size in communities experiencing warming seems to be the most common and easiest to observe organismal (at least, ectothermic) response on a global scale (Daufresne et al. 2009). It is then important to understand the conditions under which this plasticity is performed, how repeatable this response is, how variable it is between different organisms, and the reasons for this possible variability. We provide answers or suggestions to all these questions and show that it is worthwhile to study TSR not only in the context of climate change, but also in the context of the underlying basic evolutionary mechanisms responsible for the plastic response of organisms to the environment. We show that body size plasticity and canalization are condition-dependent and that these conditions are tightly linked to organismal fitness. This finding may shed light on the costs of plasticity (DeWitt 1998), and the next step should be to relate body size changes at optimal and suboptimal temperatures to bioenergetics. We also showed that there may be a tendency for specialists to be more vulnerable to global temperature changes than generalists, due to their potentially weaker ability of plastic body size response. Finally, the comparison of closely related organisms made it possible to show a high degree of conservatism in the relative descriptors of thermal tolerance, as well as in plasticity across temperature. This new knowledge adds to our understanding of the mechanisms of phenotypic plasticity in general.

## Acknowledgements

The authors would like to thank Edyta Fiałkowska and Agnieszka Pajdak-Stós for their expert advice throughout the study and Ulf Bauchinger for his valuable comments on the previous version of the text. This work was supported by the National Science Centre of Poland (2019/35/B/NZ8/03245).

## Data availability statement

All data and R code of analyses conducted are available from https://uj.rodbuk.pl/dataset.xhtml?persistentId=doi:10.57903/UJ/WUR1E5

