## Supplementary materials for "The body size and fitness match and its variability in plastic response to temperature"

**Table S1.** Parameters obtained for four models used to estimate thermal performance curves for population growth rate, proceeded following Padfield et al. (2021). rmax – maximal population growth rate, $T_{min}$ – temperature at which population growth rate switches to negative values, $T_{opt}$ – temperature at which population growth rate I maximal

| experimental population | model used | r max | Tmin | Topt |
|---|---|---|---|---|
| Int | briere2 1999() | 0.99 | 10.9 | 27.7 |
| Int | joehnk 2008() | 1.02 | 10.1 | 28.5 |
| Int | kamykowski 1985() | 0.97 | 10.1 | 28.2 |
| Int | lactin2 1995() | 1.02 | 10.1 | 28.5 |
| Un | briere2 1999() | 0.8 | 12 | 27.4 |
| Un | joehnk 2008() | 0.82 | 11.6 | 28.2 |
| Un | kamykowski 1985() | 0.79 | 11.5 | 28 |
| Un | lactin2 1995() | 0.81 | 11.6 | 28 |
| Warm1 | briere2 1999() | 0.9 | 10.6 | 27.8 |
| Warm1 | joehnk 2008() | 0.92 | 10.1 | 28.4 |
| Warm1 | kamykowski 1985() | 0.9 | 10.5 | 28.2 |
| Warm1 | lactin2 1995() | 0.91 | 10.4 | 28.2 |
| Warm2 | briere2 1999() | 1.03 | 11.8 | 28.6 |
| Warm2 | joehnk 2008() | 1.02 | 11.3 | 28.8 |
| Warm2 | kamykowski 1985() | 1.02 | 11.3 | 28.9 |
| Warm2 | lactin2 1995() | 1.05 | 11.4 | 28.9 |
| Cold | briere2 1999() | 0.83 | 9.1 | 28.8 |
| Cold | joehnk 2008() | 0.87 | 8 | 29.1 |
| Cold | kamykowski 1985() | 0.83 | 9 | 29 |
| Cold | lactin2 1995() | 0.84 | 8.9 | 29 |
| TH | briere2 1999() | 1.35 | 9.8 | 32.8 |
| TH | joehnk 2008() | 1.31 | 6.8 | 30.5 |
| TH | kamykowski 1985() | 1.25 | 10.5 | 32 |
| TH | lactin2 1995() | 1.26 | 10.3 | 31.1 |
| TNH | briere2 1999() | 1.06 | 12.1 | 30.6 |
| TNH | joehnk 2008() | 1.11 | 11.7 | 29.9 |
| TNH | kamykowski 1985() | 1.03 | 11.7 | 30.1 |
| TNH | lactin2 1995() | 1.06 | 11.8 | 29.9 |
| TN | briere2 1999() | 1.12 | 11 | 29.6 |
| TN | joehnk 2008() | 1.15 | 10.6 | 29.4 |
| TN | kamykowski 1985() | 1.11 | 11.3 | 29.6 |
| TN | lactin2 1995() | 1.13 | 11.1 | 29.5 |

**Fig. S1.** Population growth rate r obtained for stage I (black) and stage II (white) for all experimental populations. The red circles show those results that were excluded from further analyses.

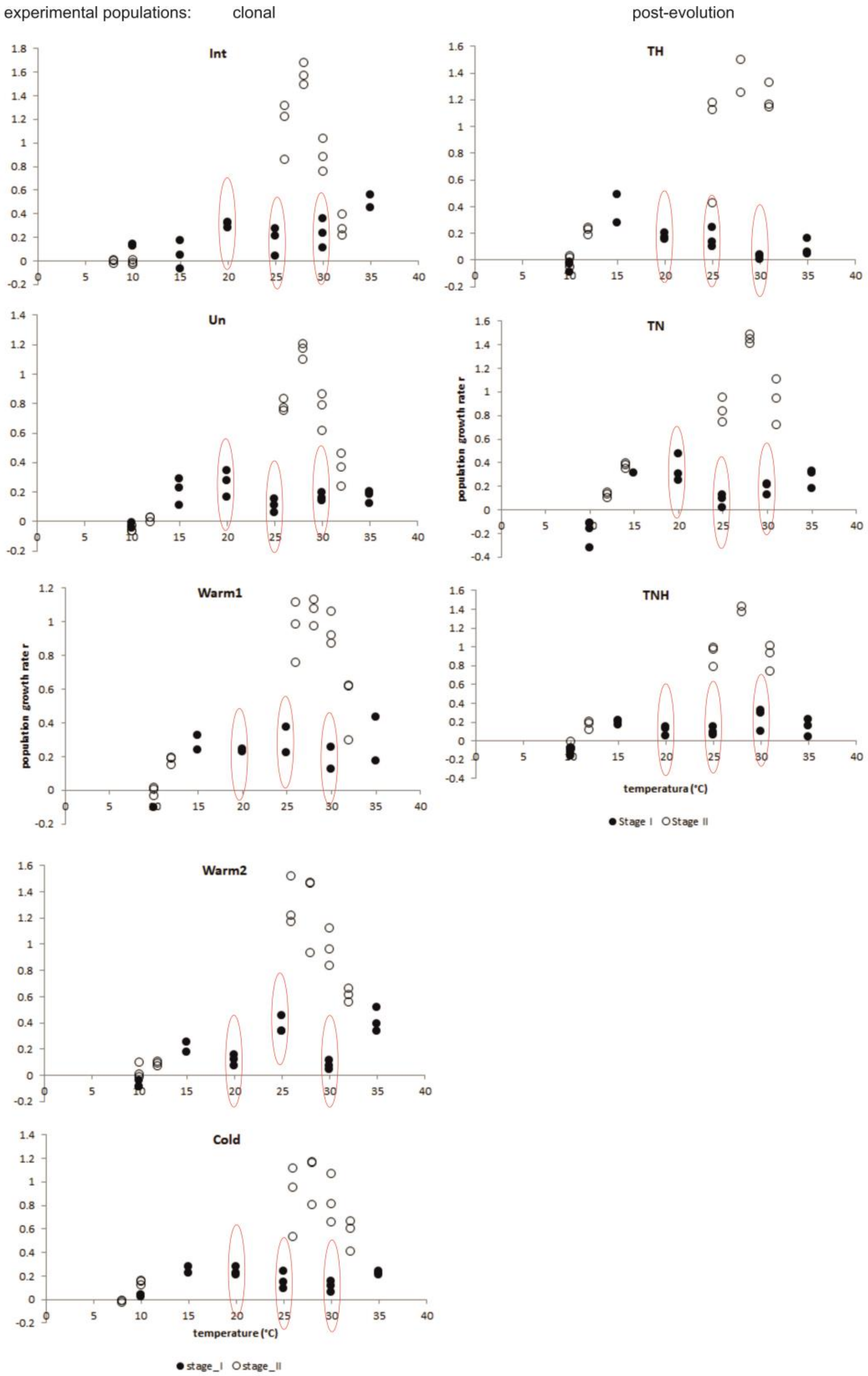

**Table S2**. Results of the statistical analyses for the body size difference between two temperatures for which the $T_{min}$ was estimated (one-way ANOVA) and between three (PEGs) or four (CGs) temperatures for which $T_{opt}$ was estimated (ANOVA with post-hoc Tukey test). CGs – clonal experimental populations, PEGs – post-evolution multiclonal experimental populations

| | $T_{min}$ estimation | | | $T_{opt}$ estimation | | | | | | | | |
|---|---|---|---|---|---|---|---|---|---|---|---|---|
| | **temp (°C)** | **F** | **p** | **temp (°C)** | **estimate** | **p** | **temp (°C)** | **estimate** | **p** | **temp (°C)** | **estimate** | **p** |
| **CGs** | | | | | | | | | | | | |
| Int | 8-10°C | 15 | 0.0002 | 26-28°C | -172.2 | 0.0271 | 28-30°C | 251.6 | 0.0003 | 30-32°C | 215.2 | 0.0036 |
| Un | 10-12°C | 15.98 | 0.0001 | 26-28°C | -135.1 | 0.0835 | 28-30°C | 199.8 | 0.003 | 30-32°C | 27.9 | 0.9644 |
| Warm1 | 10-12°C | 0.1 | 0.757 | 26-28°C | 41.1 | 0.9122 | 28-30°C | -232.7 | 0.0013 | 30-32°C | 261.5 | 0.0002 |
| Warm2 | 10-12°C | 7.22 | 0.0082 | 26-28°C | 35.9 | 0.9342 | 28-30°C | -10.4 | 0.9984 | 30-32°C | 697.6 | <0.0001 |
| Cold | 8-10°C | 7.59 | 0.0068 | 26-28°C | -460 | <0.0001 | 28-30°C | 275 | <0.0001 | 30-32°C | 318 | <0.0001 |
| **PEGs** | | | | | | | | | | | | |
| TNH | 10-12°C | 4.93 | 0.029 | 25-28°C | 177 | 0.0147 | 28-31°C | 135 | 0.0887 | | | |
| TN | 12-14°C | 16.2 | <0.0001 | 25-28°C | 218 | 0.0012 | 28-31°C | 419 | 0.0029 | | | |
| TH | 10-12°C | 15.43 | 0.0001 | 25-28°C | 454.9 | <0.0001 | 28-31°C | -13.7 | 0.9728 | | | |
